# The role of memory and perspective shifts in systematic biases during object location estimation

**DOI:** 10.1101/2021.05.29.446288

**Authors:** Vladislava Segen, Giorgio Colombo, Marios Avraamides, Timothy Slattery, Jan M. Wiener

**Affiliations:** Aging and Dementia Research Centre, Bournemouth University, UK; Department of Psychology, Bournemouth University, UK; German Center for Neurodegenerative Diseases, Magdeburg, Germany; ETH Zurich, Future Health Technologies, Singapore-ETH Centre, Singapore; Department of Psychology, Nicosia, University of Cyprus; CYENS Center of Excellence, Nicosia, Cyprus

## Abstract

Our previous research highlighted a systematic bias in a spatial memory task, with participants correctly detecting object movements in the same direction as the perspective shift, whilst misjudging the direction of object movements if those were in the opposite direction to the perspective shift. The aim of the current study was to investigate if the introduction of perspective shifts results in systematic biases in object location estimations. To do so, we asked participants to encode the position of an object in a virtual room and to then estimate the object’s position following a perspective shift. In addition, by manipulating memory load (perception and memory condition) we investigated if the bias in object position estimates results from systematic distortions introduced in spatial memory. Overall, our results show that participants make systematic errors in estimating object positions in the same direction as the perspective shift. This bias was present in both the memory and the perception condition. We propose that the systematic bias in the same direction as the perspective shift is driven by difficulties in understanding the perspective shifts that may lead participants to use an egocentric representation of object positions as an anchor when estimating the object location following a perspective shift, thereby giving rise to a systematic shift in errors in the same direction as the perspective shift.

## Introduction

Successful orientation and navigation critically depend on our ability to formulate precise spatial representations of landmarks or objects and their locations (Epstein, Harris, Stanley & Kanwisher, 1999; Postma, Kessels & van Asselen, 2004). In the lab, memory for object locations is typically assessed with tasks in which participants first encode an array of objects or environmental features from one perspective and are then asked to indicate whether the array has changed when presented from a different perspective (Diwadkar & McNamara, 1997; Schmidt et al., 2007; Hartley et al., 2007; Sulpizio, Committeri, Lambrey, Berthoz, & Galati, 2013; Montefinese, Sulpizio, Galati & Committeri, 2015; Muffato, Hilton, Meneghetti, De Beni & Wiener, 2019; Hilton, Muffato, Slattery Miellet & Wiener, 2020; Segen, Avraamides, Slattery & Wiener, 2021a; Segen, Avraamides, Slattery & Wiener, 2021b). Most previous studies employing such paradigms focused on the ability to remember object locations rather than on assessing the precision of the underlying representations. However, spatial representations can greatly vary in terms of the precision with which they are encoded (Evensmoen et al., 2013). For example, you can remember that the car is parked in a car park, or you can formulate a more precise representation in which you remember the row in which the car is parked and the relative position in this row (back, centre, front).

In our previous work (Segen, Colombo, Avraamides, Slattery & Wiener, 2021c) we designed a novel task to investigate the precision of spatial representations. The task required participants to memorise the position of an object within a room. At test, the scene would be presented from a different perspective, the object would be displaced to either the left or the right, and participants needed to decide in which direction the object had moved. To evaluate the precision of the object location representations we adopted a psychophysics approach and systematically manipulated the object displacement distances with the aim of identifying the distance at which participants would be able to reliably detect the direction of movement. Unexpectedly, we found a systematic bias that was associated with the combination of the directions of the perspective shift and object movement, which we termed Reversed Congruency Effect. Specifically, when the direction of the perspective shift and the object movement were congruent (e.g. the object moves to the right and the perspective shift is to the right) participants consistently misjudged the direction of the object movement for small object displacement distances. The opposite pattern was found in trials where the direction of the perspective shift and the object movement were incongruent (i.e. the perspective shift was in the opposite direction to the object movement direction). In this case, participants correctly identified the displacement direction regardless of the distance by which the object was moved.

Our conjecture is that the Reversed Congruency Effect is driven by biases introduced during perspective taking, with participants “dragging” the object in the same direction as the perspective shift (Figure 1). Thus, when the object remains stationary, participants would “perceive ‘‘that the object as having moved in the opposite direction of the perspective shift. Together with the actual object movement, this expectation that the object “moves” in the same direction as the perspective shift would yield the observed Reversed Congruency Effect. Specifically, if the object moved in the opposite direction to the perspective shift, participants would perceive the object movement to be larger due to the expectation that the object follows the perspective shift. Whilst, in situations when the object moves in the same direction as the perspective shift, participants may incorrectly perceive the object movement direction, as the change in the object position may not be large enough to overcome their expectation regarding the new object position following a perspective shift. Yet, in trials when object movement was large enough, the effect of the perspective shift related expectation of object movement is overcome allowing participants to correctly detect the direction in which the object moved.

**Figure 1.**
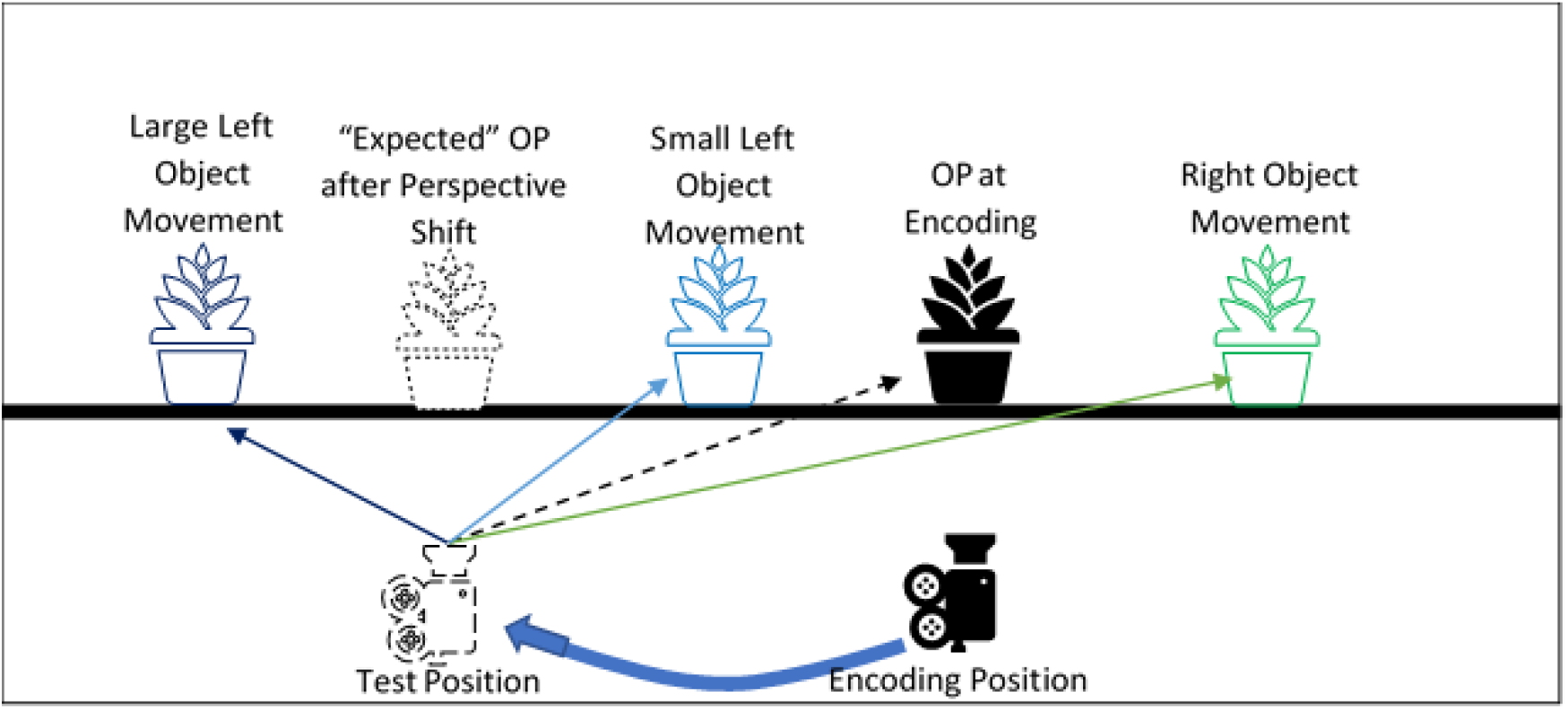
Schematic of the Reversed Congruency Effect: The black plant and camera represent the position of the object (OP) and camera at encoding. The dotted camera represents the position at test following a perspective shift to the left. The dotted plant represents the “expected” position of the object following a perspective shift if participants “drag” the object with them. Given the new position (dotted camera) it appears that even if the object was stationary (black plant) that the object has moved right i.e. perspective shift induced object motion. The green plant represents small movement to the right, which is perceived to be much larger due to the perspective shift induced object motion. Whilst small left movements (light blue plant) are perceived as right movements due to being further to the right than the “expected” object position, yet, when the movements to the left (congruent with the direction of the perspective shift) were large enough (i.e. dark blue plant) participants could correctly detect the movement direction.

Although this explanation is in line with our empirical data, our original study (Segen et al., 2021c) did not allow us to directly investigate if the Reversed Congruency Effect described above was primarily driven by the proposed perspective shift related bias in which participants drag the object in the same direction as the perspective shift. Alternatively, it is possible that the Reversed Congruency Effect relied on the presence of the object in both the encoding and test phase and that the comparison of the object locations across those stimuli gave rise to the observed bias.

Following up on our previous work, the first aim of the current study was to investigate whether perspective shifts lead to a systematic bias in the remembered object positions. This question is particularly important also because many studies investigating spatial memory and perspective taking ability present static images across different perspectives and could be subject to the same effect (Diwadkar & McNamara, 1997; Schmidt et al., 2007; Hartley et al., 2007; Sulpizio et al., 2013; Montefinese et al., 2015; Muffato et al., 2019; Hilton et al., 2020; Segen et al., 2021a; Segen et al., 2021b). To address this question, we designed a task in which participants first encoded the position of an object. Then, they were presented with an image of the same scene but from a different perspective but without the object and had to indicate the position of the object. If, as argued above, the Reversed Congruency Effect was driven by a perspective shift related bias, we expect that participants will produce systematic errors in the same direction as the perspective shift. That is, if the perspective shift is to the left, participants would place the object further to the left of its actual position.

Our second aim was to investigate whether the potential perspective shift-related bias is related to memory processes. It is well known that spatial memory is prone to distortions. For example, when drawing sketch maps of environments from memory, participants often draw non-orthogonal junctions as 90° junctions and straighten the curved street segments (Wang & Schwering, 2009). In addition, distance estimates are influenced by the presence of physical or geographical borders (Uttal, Friedman, Liu & Warren, 2010). Memories for object locations are also prone to systematic biases. That is, many studies have shown that object location estimates tend to “move” towards category prototypes (Huttenlocher, Newcombe, & Sandberg, 1994; Crawford & Duffy, 2010; Holden, Curby, Newcombe & Shipley, 2010; Huttenlocher, Hedges & Duncan, 1991). For example, when asked to memorise the location of a dot in a circle, participants divide the circle into quadrants and estimate the dot position closer to the centre of each quadrant (Huttenlocher et al., 1991).

Additionally, previous research suggests that spatial perspective taking is differently affected depending on whether the task needs to be solved by relying on spatial memory. Specifically, Hartley et al. (2007) showed that reliance on spatial memory leads to greater difficulties in spatial perspective taking. The authors suggested that this can be explained by the need to manipulate the whole scene to achieve perspective-taking if the representation is held in memory. In contrast, when participants can see the scenes from both perspectives simultaneously (perception condition) it is possible to use piecemeal rotation of each element in the scene to ensure that the positions between the two scenes match. Following this explanation, we would expect that the perspective-shift related bias would only be apparent in the memory condition, where perspective taking itself may be more complex.

We investigated whether memory contributes to the predicted perspective shift related bias in the object locations by creating two conditions: in the memory condition, participants first saw the image of a scene with the target object during encoding, and, following a short delay, the second image showing the same scene from a different perspective but without the object. Their task was to indicate, on the second image, the position of the object. In the perception condition, participants performed the same task but the two images were presented simultaneously on two adjoining computer screens. If memory contributes to the systematic bias introduced by the presence of a perspective shift, we expect a stronger bias in the memory condition than in the perception condition. However, if the effect is driven by the introduction of the perspective shift and is independent of memory, we expected similar biases in the two conditions.

## Method

### Participants

Seventy-seven participants took part in the experiment (Mean age=19.94 years, SD =2.35; age range = 18–32 years; 49 females and 28 males) with thirty-nine participants completing the Memory condition and thirty-eight the Perception condition. Participants were recruited through Bournemouth University’s participant recruitment system and received course credit for their participation. All participants gave their written informed consent in accordance with the Declaration of Helsinki (World Medical Association, 2013).

### Materials

#### Virtual environment

The virtual environment was designed with 3DS Max 2018 (Autodesk Inc) and consisted of a square room (9.8 m x 9.8 m) that contained famous and easily recognisable landmarks on its walls (Hamburger & Roser, 2014). A teal plank was placed diagonally in the middle of the room (14 m long). During encoding, an object was placed on that plank at one of 18 predefined positions that were 14, 28, 42, 84, 98, 112, 168, 182 and 192 cm to the left or to the right of the centre of the plank. The object was removed during testing and 37 markers appeared on the plank serving as possible response locations (Figure 2B).

**Figure 2:**
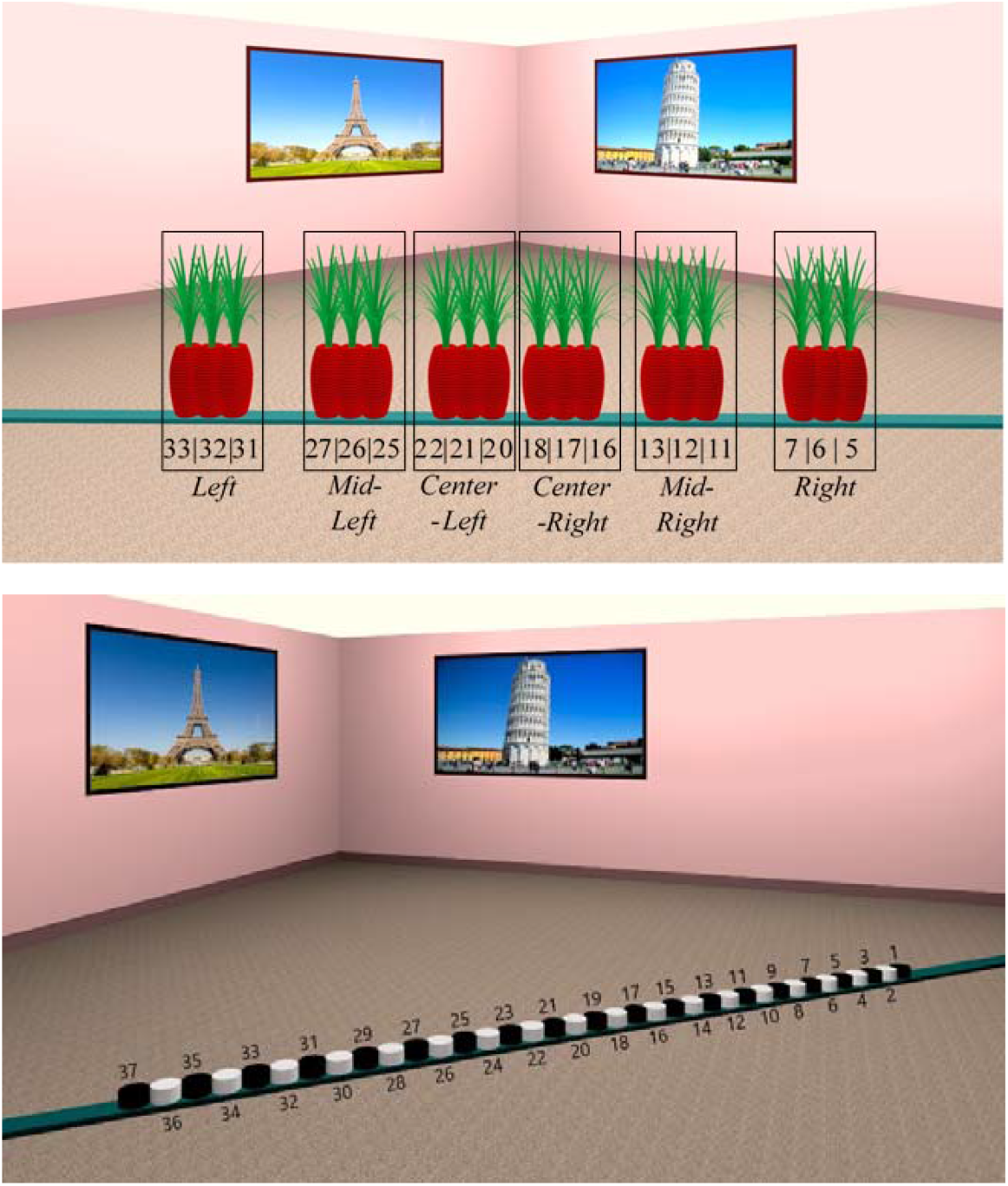
A: Example stimuli superimposing all of the possible object positions ranging between 5 to 33 (positional markers in Figure 2B) and the corresponding six Clusters (Left, Mid-Left, Center-Left, Center-Right, Mid-Right, Right); B: Example of Test stimuli containing the positional markers from 1 to 37 that participants needed to select to estimate object position

To analyse participant responses, we created six groups containing the three object positions (*Left, Mid-Left, Center-Left, Center-Right, Mid-Right, Right*) that were close to each other, i.e. objects positions at 14, 28 and 42 cm to the left of the centre were grouped together (Figure 2A). From hereon we will refer to those object groups as Clusters.

The visual stimuli were presented on a 40-inch screen at a resolution of 1920×1080px and subtended 47.7° x 28° at a viewing distance of 1 meter. The experimental stimuli were renderings of the environment with a 60° horizontal field of view (FOV), a custom asymmetric viewing frustum that resembles natural vision with a 15% shift in the vertical field of view was used (Franz, 2005; Figure 3).

**Figure 3.**
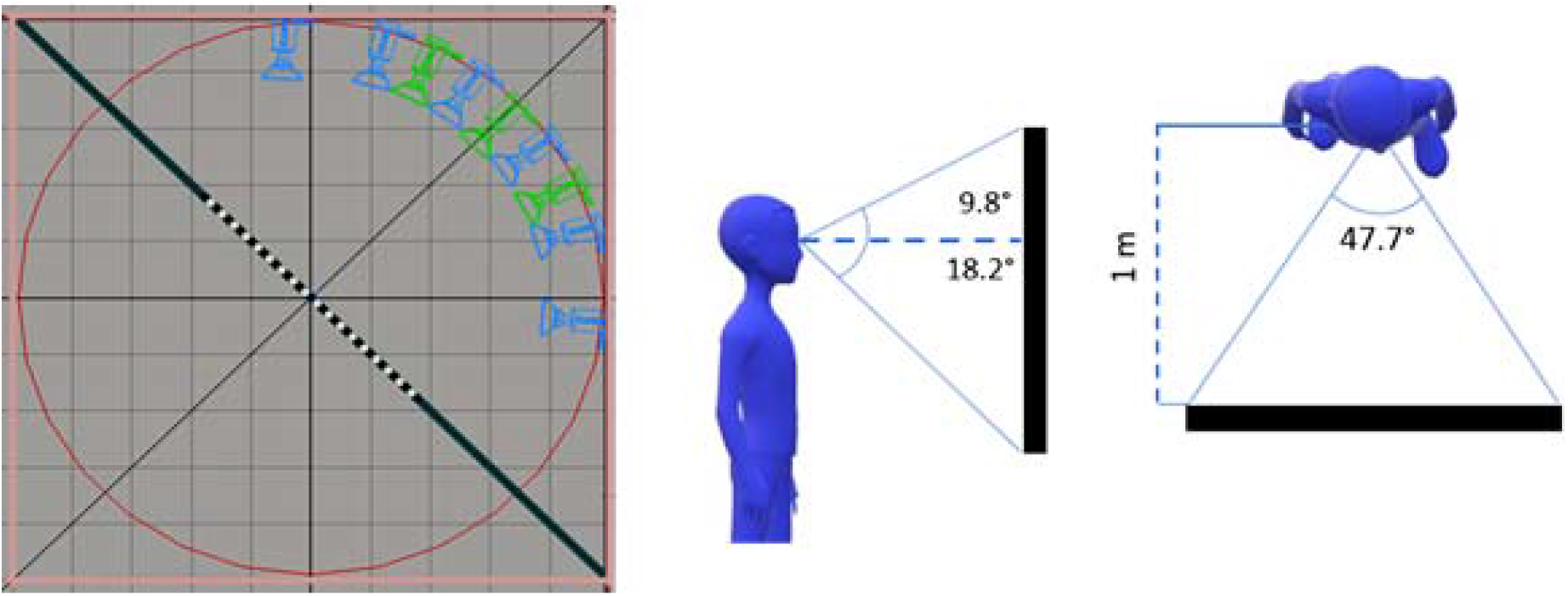
Left Schematic of encoding (green) and test (blue) camera positions arranged in an invisible circle in the environment; Right A representation of how participant position related to the stimulus display.

The cameras were arranged in an invisible circle around an invisible diagonal line that was perpendicular to the plank. The encoding stimuli were rendered from three possible camera positions (Figure 3). The test stimuli were rendered from a different viewpoint with a 30° perspective shift either to the left or to the right of the encoding viewpoint. In both encoding and test stimuli, the room corner and one poster at each side of the corner were visible.

Stimuli were presented with OpenSesame 3.1.7 (Mathôt, Schreij, & Theeuwes, 2012). In the Memory condition, the stimuli were presented on a single monitor and in the Perception condition stimuli were presented across two monitors. Responses were made with a standard keyboard that was labelled such that a different key corresponded to each of the 37 possible positional markers. Participants had to choose the marker that they thought corresponded to the position of the object during encoding, and to press the key that corresponded to that marker (Figure 2B).

##### Procedure

Each experimental trial started with the presentation of an instruction prompting participants to remember the location of the object (750 msec). This was followed by a display containing a fixation cross and a scrambled stimuli mask (500 msec). In the Memory condition, this was followed by the encoding phase, in which participants were presented for 5 seconds with an image of the scene that depicted the object in one of the 18 possible positions in the room, taken from one of three camera positions. After the encoding phase, participants were again presented with a fixation cross and a scrambled stimuli mask for 500 msec. In the test phase that followed, they were presented with another image that was taken after a 30° perspective shift. In this picture, the object was removed, and 37 labelled markers appeared on the plank which participants used to indicate object locations (Figure 4A). In the perception condition, participants were presented with the encoding and test stimuli simultaneously across two screens (Figure 4B). In both conditions, participants were free to take as long as they needed to make a response.

**Figure 4.**
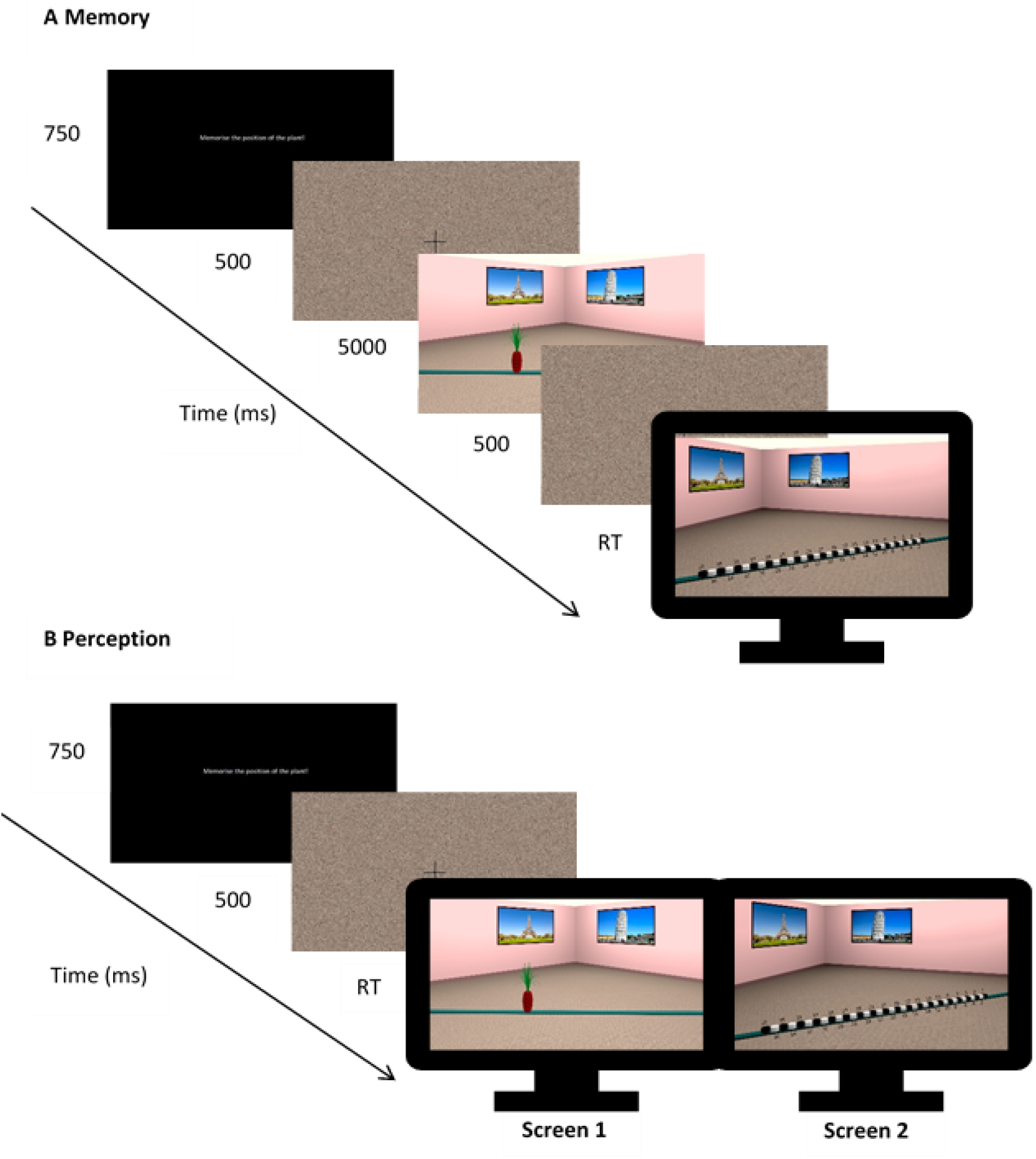
Trial structure in the Memory (A) and Perception (B) conditions

### Design

A between-subject design was adopted, and block randomization was used to assign participants to the Memory or Perception condition. This ensured an approximately equal number of participants in each condition. Overall, the experiment included 108 experimental trials presented in randomised order with the experiment taking on average about 30 minutes.

### Data Analysis

Statistical analyses were carried out using R (R Core Team, 2013). Data were analysed with linear mixed-effects models (LME) using LME4 (Bates et al. 2015) in R (R Core Team, 2013). Effect coding was used as contrasts for fixed factors, which were all categorical variables. The absolute error model included the by-item intercepts as well as a by-subject intercept and slope for Perspective Shift Direction (PSD). Prior to analysis, outlier responses were removed using the interquartile range method on individual absolute error (cm) distributions, which led to a total 3.3% data loss.

## Results

### Absolute error

We first examined the effect of Condition (*Memory* vs *Perception*), Cluster (*Left, Mid-left, Centre-left, Center-right, Mid-right and Righ*t) as well as the Perspective Shift Direction (PSD; *Right* vs *Left*) on absolute error (cm) (full results reported in supplementary materials). Interestingly, the results show that the absolute error was higher in the Perception compared to the Memory condition (β=3.002, SE=1.052, t=2.854) and there were no main effects of Cluster or Perspective Shift Direction. An interaction was found between Condition and Cluster, such that in the *Memory* condition errors in the *Left* cluster were lower than in the *Perception* condition (β=1.862, SE=0.393, t=4.736), no reliable differences between conditions was found for any of the other clusters. In addition, we found an interaction between Cluster and PSD, with higher errors in the *Right* cluster (β=3.271, SE=0.894, t=3.658) and lower errors in the *Left* cluster (β=-2.459, SE=0.895, t=-2.747) when the PSD was to the *Left*. This suggests that errors increased when perspective shifts resulted in movements away from the object cluster. This effect was amplified in the *Perception* compared to the *Memory* condition, with an even greater increase in errors in the *Right* (β=0.813, SE=0.393, t=2.068) and *Mid-right* (β=1.529, SE=0.395, t=3.870) cluster when the perspective shifted to the *Left* in the *Perception* condition and a greater decrease in errors in the *Center-Left* (β=-0.969, SE=0.393, t=-2.463) and *Mid-Left* (β=-1.936, SE=0.394, t=-4.918) clusters.

### Signed Errors

We did not find differences in absolute errors as a function of PSD (*Left* and *Right*) and we have no reason to assume that perspective shifts to the left or the right would be qualitatively different. Errors to the left had a negative sign (i.e. −30 cm) and errors to the right had a positive sign (i.e. 30cm). However, since we are primarily interested in the direction of the errors as a function of the direction of the perspective shift, we,, therefore, multiplied (folded) all of the errors where the perspective shifted to the left by −1. Following the folding procedure, the resultant positive errors indicate errors in the same direction as the perspective shift (i.e. perspective shift is to the left and the errors are to the left) and negative errors indicate errors in the opposite direction (i.e. perspective shift is to the left and the errors are to the right). An LMM with Condition as a fixed effect revealed that overall, errors were positive (Intercept: β=10.927, SE=2.013, t=5.429). In other words, participant responses were biased towards the direction of the perspective shift (Figure 5). Signed errors did not differ between the *Memory* and the *Perception* conditions (β=-0.672, SE=1.551, t=-0.433).

**Figure 5.**
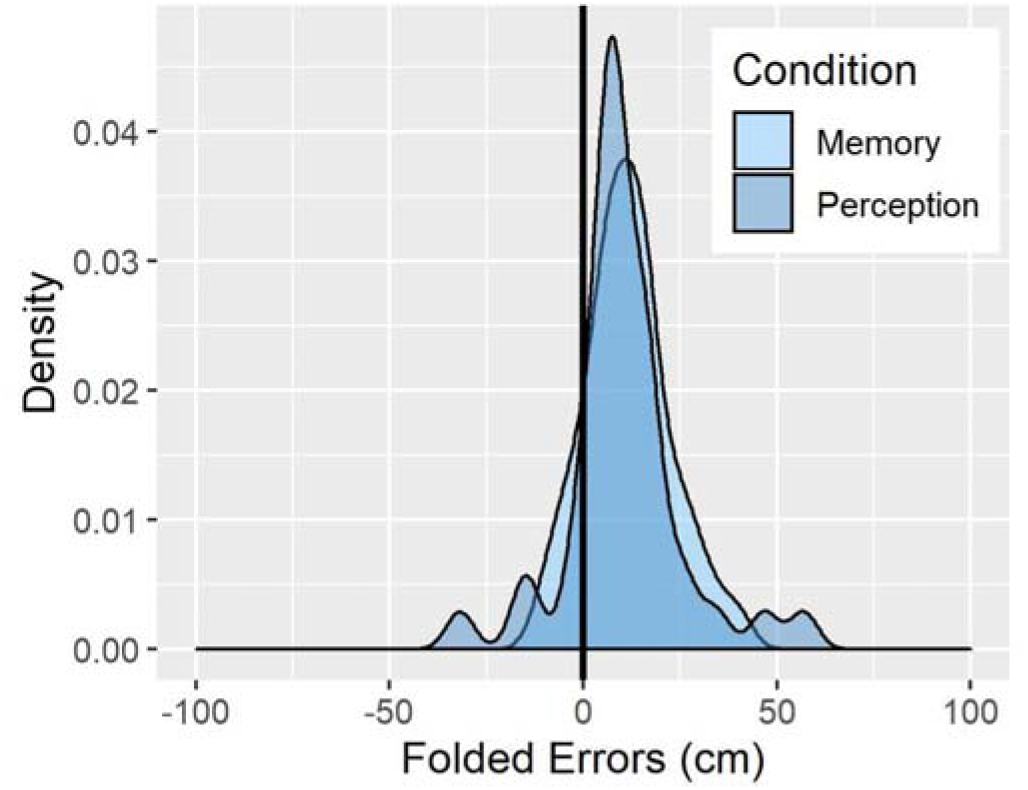
Density plot of Signed Errors (cm) across the Memory and Perception conditions

### Role of Object Position

Given previous reports of systematic biases in object location memory (Huttenlocher et al., 1991) towards a “category” prototype, we examined if object positions had an impact on participants’ errors. To do so we calculated, using the response markers, the range of responses for each of the 18 object positions, such that the value of 0 corresponds to responses in which the participants placed the object in the correct position, negative values represent errors made to the left, and positive values indicate errors to the right. Figure 7 displays histograms of responses for each object position. To investigate if participants’ responses for each object position were significantly different from zero, thus indicating a systematic bias, we ran one-sample t-tests for each object position separately for the Memory and Perception conditions.

As it is not clear what prototypes participants might have used in the current task, we evaluated different alternatives suggested by the previous literature. For example, one possibility is that participants remembered objects to be closer to the centre of the screen (conceptually similar to central tendency bias [Allred, Crawford, Duffy & Smith, 2015], Figure 6A). If participants indeed used the centre of the screen as the prototypical object position, we would expect them to make errors to the left for object positions 5 to 18, and to the right for object positions 20 to 33 (Figure 6A). Another possibility is that participants divided stimuli into the left and right half and used the centre of each half as prototypical locations (Huttenlocher, et al., 1994; Crawford & Duffy, 2010). If participants used the centre of those halves as prototypes we would expect a leftward bias in object positions 5-7 and a rightward bias for object positions 11 to 18 as this would bring objects positioned on the right closer to the centre of the right half of the plank. For the left half of the stimuli we would expect a leftward bias for object positions 20 to 27 and a rightward bias for object positions 31 to 33 (Figure 6B). Another possibility is that participants used more fine-grained categories in which the object in the centre of each of the six object clusters functioned as a category prototype (Figure 6C; Holden et al., 2010). This way, in the cluster consisting of object positions 31,32, and 33, participants would estimate the object positions to be closer to object position 32.

**Figure 6:**
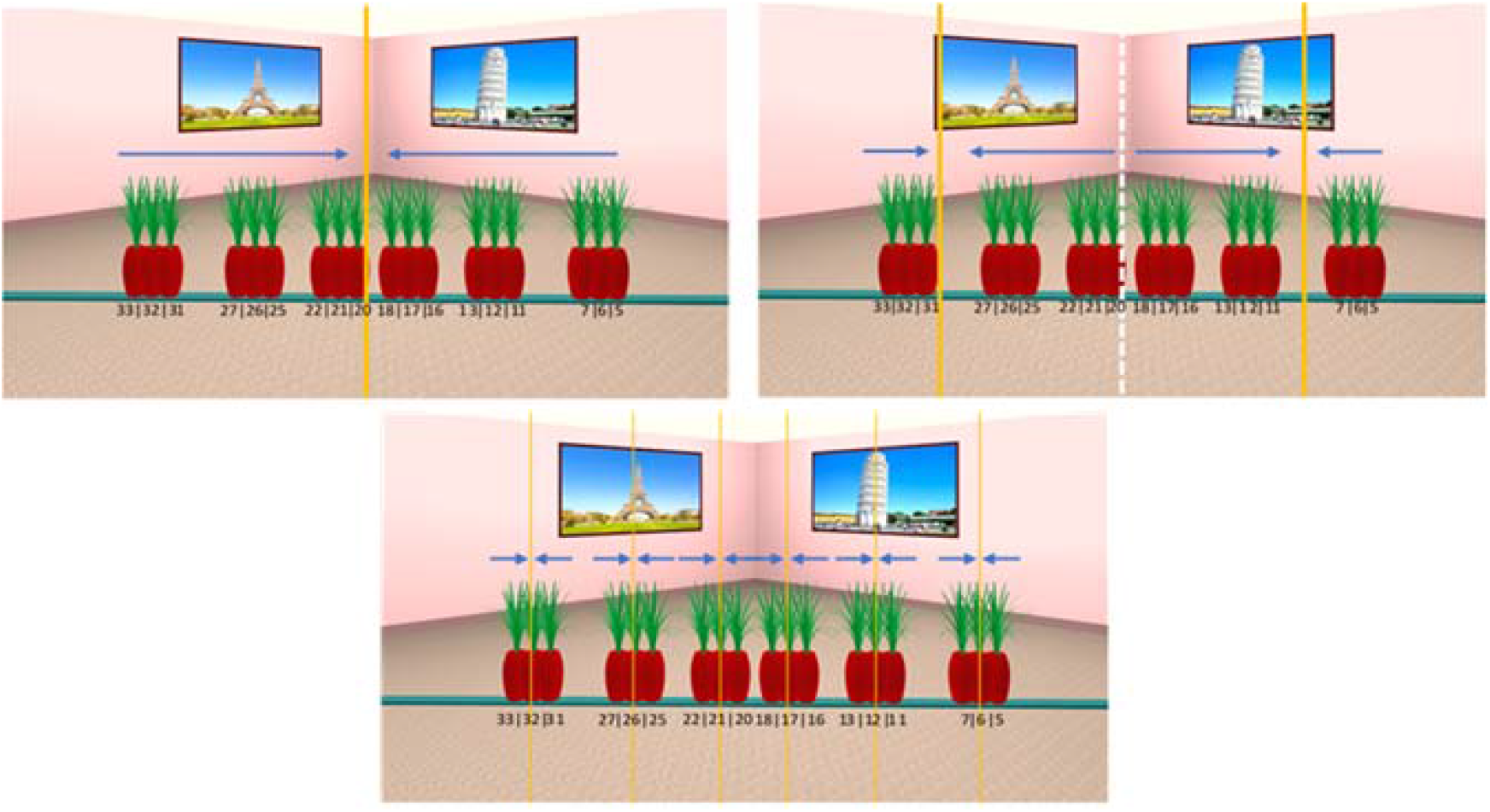
Examples of possible object location prototypes that participants may use with the blue arrows indicating the expected bias direction. Orange lines indicate prototype locations. (A) Center of the screen, (B) center of the left and right side of the screen or (C) center of the cluster used as a category prototype.

Our results showed that for objects positioned at the extremes of the possible object positions (most leftward [i.e. 33,32,31] and most rightward [5, 6, 7] positions), participants made errors away from the extreme values (the positional markers on both ends) (Figure 7). For example, for object positions 33 and 32 which are on the left side of the plank, participants made more errors to the right, whilst for objects positions 5, 6 and 7 that are on the right, participants made more errors to the left. This result is partly in line with the category prototypes depicted in Figure 6A and 6B. However, for the more central object positions, we found a slight bias to the right that is not consistent with any of the possibilities we described (Figure 6).

**Figure 7:**
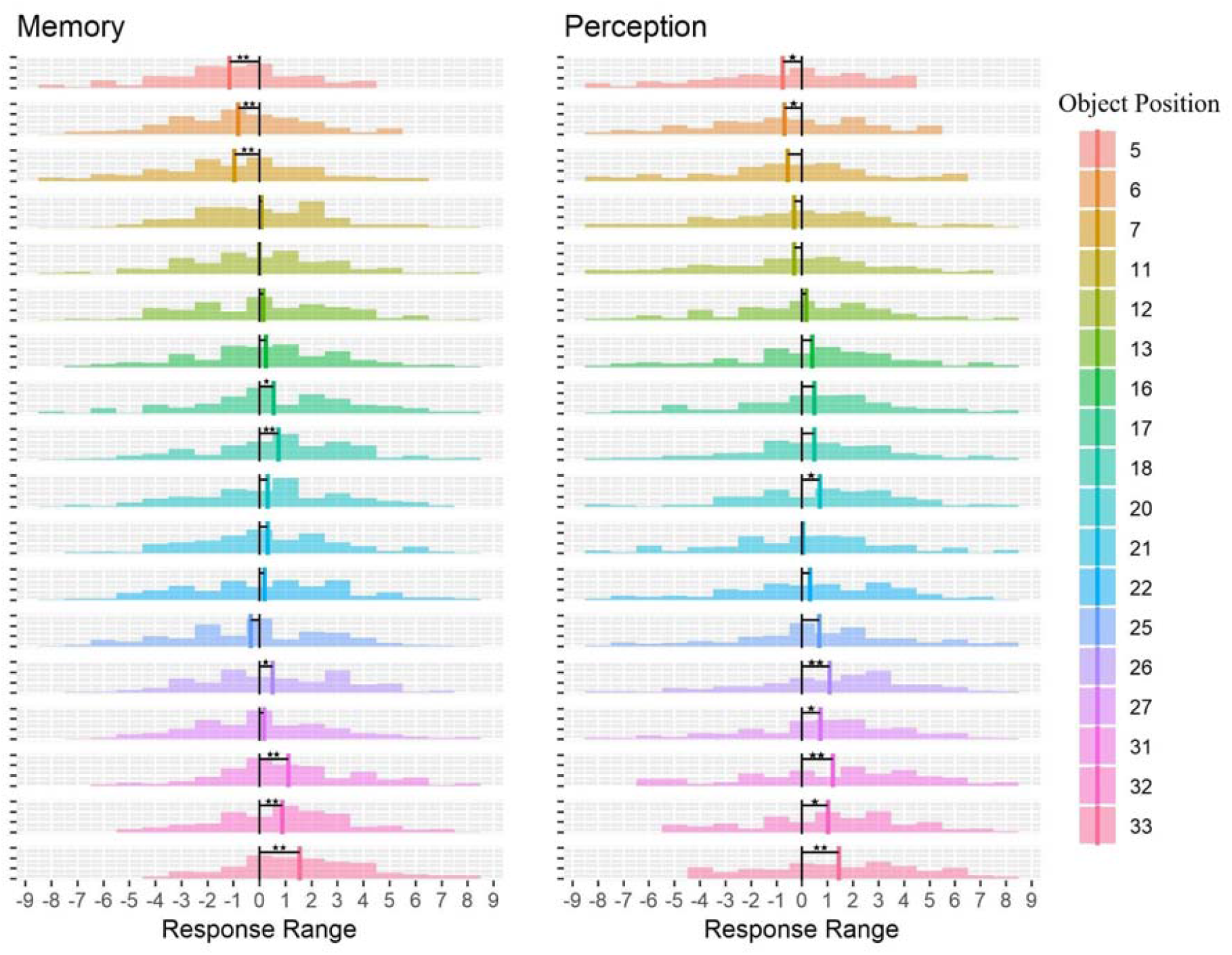
Distribution of the response range for each object position as a function of Condition (Memory and Perception)

We have also looked at directional errors with the complete model reported in the supplementary materials. As reported above, the direction of the perspective shift determined the direction of the errors. That is, if the perspective shift was to the right then the errors were to the right as well (positive errors). This was the case across all but the most leftward and rightward clusters, for which we found that participants made errors away from the extremes such that the direction of the perspective shift no longer determined the direction of the errors. Instead, participants made more errors to the right in the left cluster, with the opposite pattern of errors found for the most rightward cluster (Figure 8).

**Figure 8:**
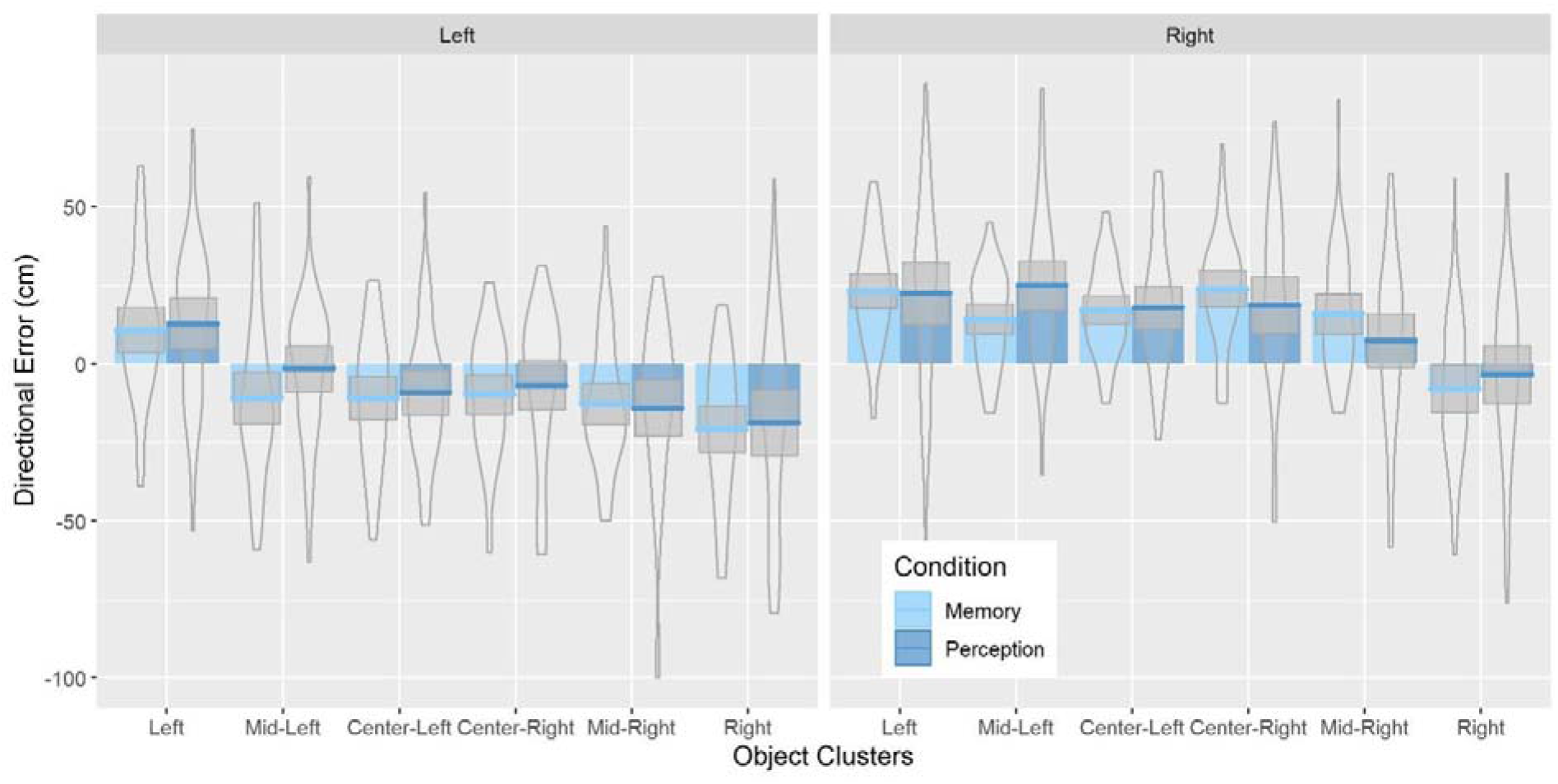
Bar plots for directional errors as a function of Perspective Shift Direction, Condition and Object Clusters with mean (solid line) and 95% CIs (grey shaded area) with violin plots behind

## Discussion

The aims of the current study were twofold: the first aim was to investigate if perspective shifts systematically bias estimates for object positions. The second aim was to investigate if the proposed bias in object position estimates arises from distortions in spatial memory. To do so, we explored error patterns in a task in which participants estimated the position of an object following a perspective shift either with or without a memory delay. Consistent with our expectations, we found that participants’ errors were systematically biased in the direction of the perspective shift, we termed this as the *perspective shift related bias*. Importantly, this *perspective shift related bias* was observed in both the *Memory* and *Perception* conditions, suggesting that it is not related to systematic distortions in memory.

But how can this systematic *perspective shift related bia*s in object location estimation be explained? Spatial perspective taking can be achieved either by relying on an allocentric representation or by mentally transforming an egocentric representation (King et al, 2002; Hegarty & Waller, 2004). Yet, if participants relied solely on an allocentric representation in which the position of the object was encoded relative to other features in the environment, their own position and movement in the environment should not influence their responses and perspective shifts should not result in systematic biases (Ekstrom, Arnold & Iaria, 2014). Thus, the presence of the *perspective shift related bias* in the estimations of object locations in the direction of the perspective shift, suggests an egocentric influence on the estimates.

Specifically, we believe that uncertainty about the exact nature of the perspective shift leads to uncertainty about the exact object location, which in turn results in participants biasing their estimates towards the encoded egocentric location of the object. This idea is conceptually similar to the anchoring and adjustment heuristic proposed by Tversky and Kahneman, (1974), which posits that, when uncertain, people make decisions/responses using an initial estimation, an anchor that they then adjust to correct for errors. Interestingly, these anchors are often based on egocentric representations (Epley, Keysar, Van Boven, Gilovich, 2004; Gilovich, Medvec, & Savitsky, 2000; Keysar, Barr, Balin, & Brauner, 2000). For example, people often use their own experience as an anchor when estimating how their actions affect others (Gilovich et al., 2000) and when making judgements about how others perceive ambiguous stimuli (Epley et al., 2004). In the current task, participants may have used the original egocentric relation of self to object as an anchor, which would result in participants dragging the object with them following a perspective shift. Adjustments are then made, taking into account the available information about the perspective shift, i.e. changes in the position of other features in the environment. However, if participants are uncertain about the exact nature of the perspective shifts, these adjustments are not sufficient, resulting in estimates that are biased towards the anchor (Tversky & Kahneman, 1974; Quattrone, 1982). This leads to a systematic shift in object position estimates in the direction of the perspective shift giving rise to the *perspective shift related bias*.

We also found that when the perspective shift increased the distance to the object, participants were less accurate in estimating its position and displayed a larger *perspective shift related bias*. This pattern flipped in situations when the distance to objects decreased following a perspective shift, showing that participants were more accurate and less biased in estimating object positions when they were closer to them. One potential explanation for this is that there is greater compression of space for locations that are further away. Therefore, the difference between neighbouring object positions may become less pronounced the further away they are, making it harder to choose the appropriate position as the position markers are smaller and closer together. Given that the markers “appear” closer together for further away locations, it is also possible that a larger number of positional markers are considered as plausible estimates (as they are all close together), leading participants to accept positions that are further away from the actual object position but are closer to the original egocentric vector that is used as an anchor. This is in line with the idea that adjustments of the initial anchor are made until a plausible estimate is reached (Epley et al., 2004).

An alternative explanation for the *perspective shift related bias* relates to the specifics of the camera movement during the perspective shift. In our study, the camera moved on a circle such that a perspective shift to the left was realised by a camera translation to the left and a camera rotation to the right in order for the camera to remain directed towards the same point in the room. Such camera movements are typically used in spatial perspective taking tasks (Montefinese et al., 2015; Muffato et al., 2019; Hilton et al. 2020; Segen et al., 2021a; Segen et al., 2021b; Sulpizio et al., 2013). This combination of camera translation and rotation is chosen to ensure that the same part of the scene is visible in the images before and after the perspective shift. However, it produces images that can look surprisingly similar, and, as a result, may cause participants to underestimate the size of the perspective shift. Underestimation of the perspective shift may lead participants to think that the camera movement was smaller than it was, yielding a bias in responses to the direction of the perspective shift. While we cannot distinguish between this explanation and the anchoring heuristic in the current study, we recently ran a follow up experiment in which we systematically manipulated the way the camera moved during a perspective shift (Segen et al., in prep). Results from this follow-up experiment provides support for the anchoring hypothesis and suggests that the influence of camera rotations is marginal.

The second aim of this study was to investigate if the bias in object position estimates result from systematic distortions in spatial memory. Importantly, we did not find a difference in the perspective shift related bias between the memory condition and perception condition showing that the systematic bias in errors in the direction of the perspective shift is not introduced by memory. Additionally, we also found a small difference in absolute errors, with participants performing better in the memory than in the perception condition, thus further highlighting that the observed defects are unlikely to be driven by memory processes. Such findings contrast with previous research showing that biases in object location estimations are typically introduced by post-encoding processes (Crawford, Landy, & Salthouse, 2016). For example, when participants estimate city locations from memory they incorrectly place Montreal farther north than Seattle, influenced by their prior knowledge of Canada being to the north of the U.S (Friedman, Kerkman, Brown, Stea, & Cappello, 2005). In general, biases in object-location memory are typically explained by a post-encoding Bayesian combination of more uncertain fine-grained information with the more certain category knowledge (Huttenlocher et al., 1991).

Yet, given our interpretation that the systematic bias is driven by processes underlying the perception/understanding of the perspective shift, it is not entirely surprising that we do not find differences between the memory and perception conditions. It should be noted that participants need to engage in spatial perspective taking in both situations, with the only difference being that in the memory condition they need to rely on a stored representation which they should either manipulate to match the test viewpoint or use as a reference to which the test stimuli viewpoint is matched.

To further investigate the role of memory in object location estimation we focused on the positions of the objects in the environment, as object location memory has been shown to be biased towards category prototypes (i.e. centre of the screen, centre of the quadrant) (Huttenlocher et al., 1991; Crawford, Landy, & Salthouse, 2016). Consistent with the prominent models of object location memory i.e. the category adjustment model (Huttenlocher et al., 1991)/Dynamic Field Theory (Simmering, Spencer & Schöner, 2006; Spencer & Hund, 2002), we found that for the most leftward and rightward object positions, errors shifted away from the extremes towards the centre. However, we did not find a systematic shift away from the central positions towards category prototypes that would be expected based on these models. This is consistent with our findings that the systematic bias is not introduced by memory, as the bias towards a prototype is a phenomenon that relates specifically to object-location memory and increases with memory delay.

Notably, we did find a slight shift in errors to the right for the more central positions. A possible explanation for this bias is that the cameras were always directed towards the same spot in the environment that was slightly to the left of the center. If participants did not perceive this slight rotation and assumed that the camera faced the centre of the room, they may have remembered the object to be slightly to the right. However, even if this was the case, the effect is very minor and overall our results point to a systematic bias away from the extremes rather than towards a specific prototype and performance is mainly influenced by the perception/understanding of the perspective shift rather than distortions introduced in memory.

We also found that the absolute errors were lower in the memory condition than the perception condition. This was surprising as the requirement to memorise the encoding stimulus should have increased the cognitive demand which should have led to reductions in performance. However, the differences between the perception and memory condition were small and resulted from longer tails in the perception condition. We therefore believe that it is unlikely that there are fundamental differences between the memory and perception conditions.

Lastly, we turn our discussion to the relationship between the current findings of the *perspective shift related bias* and the *Reversed Congruency Effect*, which manifested itself in better performance in estimating object movements that are in the opposite direction to the perspective shift and misjudgement of smaller movements in the same direction as the perspective, that we found in our previous study (Segen et al., 2021c). The unexpected finding of the *Reversed Congruency Effect* was an important motivator for the current study as it was the first report of a systematic bias related to the direction of the perspective shift. We proposed that the *Reversed Congruency Effect* was driven by the *perspective shift related bias*. Specifically, if participants estimated the original object position to be shifted in the same direction as the perspective shift, as results from this study show, movement of an object in the opposite direction to the perspective shift would be perceived as larger and thus detected more easily. However, when the object moves in the direction of the perspective shift, the actual movement is attenuated by the expectation that the initial object position is “shifted” in the same direction. In such situations, smaller object movements may give rise to the impression of the object having moved in the opposite direction, as the expectation of original object position following a perspective shift may be shifted more in the direction of the perspective shift than the actual object movement.

Our findings of a reduced *Reversed Congruency Effect* with the use of additional information in the environment (i.e. columns that acted as environmental cues; Segen et al., 2021c) align with the anchor and adjustment explanation for the perspective shift related bias that we observe in the current study. Specifically, since adjustments are made on the basis of the information available (Northcraft & Neale, 1987; Tversky & Kahneman, 1974), and in our case this information is about the perspective shift, increasing the reliability of this information should reduce the biases introduced by the anchoring and adjustment heuristic. We contend that these additional cues result in a more precise understanding of the position of the object in space and a better understanding of the perspective shift. This reduces the uncertainty about the object position after the perspective shift and thus the weight given to the egocentric anchor while improving the adjustment process.

To conclude, the current study shows that participants make systematic errors in the same direction as the perspective shift when estimating object locations across different perspectives. This *perspective shift related bias* is present even in a perceptual version of the task and is likely driven by difficulties in understanding/perceiving the perspective shifts. We believe that the egocentric spatial relations between observer and target object acts as an anchor that participants fail to adequately adjust after the perspective shift. As a result, they make responses that are biased in the direction of the perspective shift. However, more research is needed to fully understand the mechanisms that give rise to the perspective shift driven bias in object location estimation. Importantly, the current findings are a conceptual replication of the *Reversed Congruency Effect* we reported in our previous study (Segen et al., 2021c). The presence of the perspective *shift related bias* across two different experimental paradigms (different sizes of perspective shifts, different tasks [determine direction of object movement vs estimate object positions]) suggests that this is a robust effect that may translate to other studies that rely on static stimuli and perspective shifts. Thus, it is paramount for researchers who use similar paradigms to be mindful of this bias as it can greatly influence the interpretation of their results.

## Notes

https://doi.org/10.6084/m9.figshare.14701467.v1

https://doi.org/10.6084/m9.figshare.14701461.v1

https://doi.org/10.6084/m9.figshare.14701455.v1

### Competing Interest Statement

The authors have declared no competing interest.

