## Supplementary material for "The role of memory and perspective shifts in systematic biases during object location estimation": Supplimentary Materials

### Supplementary Materials

#### Absolute error analysis

Table 1 Coefficients from Absolute Errors (cm) LMM analysis

|  | **Absolute Error** | | |
| --- | --- | --- | --- |
| *Predictors* | *Estimates* | *std. Error* | *t-value* |
| (Intercept) | 36.076 | 1.112 | **32.452** |
| Condition (*Perception*) | 3.002 | 1.052 | **2.854** |
| Cluster (*Right*) | -0.661 | 0.894 | -0.739 |
| Cluster (*Mid-right*) | 0.038 | 0.895 | 0.043 |
| Cluster (*Centre-left*) | 0.087 | 0.895 | 0.097 |
| Cluster (*Left*) | 0.327 | 0.895 | 0.365 |
| Cluster(*Mid-left*) | 0.089 | 0.895 | 0.099 |
| PSD (*Left*) | -0.391 | 0.573 | -0.682 |
| Condition (*Perception*)* Cluster (*Right*) | -0.632 | 0.393 | -1.608 |
| Condition (*Perception*)* Cluster (*Mid-right*) | 0.209 | 0.395 | 0.530 |
| Condition (*Perception*)* Cluster (*Centre-left*) | -0.244 | 0.393 | -0.620 |
| Condition (*Perception*)*Cluster (*Left*) | 1.862 | 0.393 | **4.736** |
| Condition (*Perception*)* Cluster (*Mid-left*) | 0.105 | 0.394 | 0.266 |
| Condition (*Perception*)* PSD(*Left*) | -0.486 | 0.446 | -1.089 |
| Cluster (*Right*)*PSD (*Left*) | 3.271 | 0.894 | **3.658** |
| Cluster (*Mid-right*)*PSD (*Left*) | 1.401 | 0.895 | 1.565 |
| Cluster (*Centre-left*)*PSD(*Left*) | 0.413 | 0.895 | 0.462 |
| Cluster (*Left*)*PSD (*Left*) | -2.459 | 0.895 | **-2.747** |
| Cluster (*Mid-left*)*PSD(*Left*) | -0.452 | 0.895 | -0.505 |
| Condition (*Perception*)* Cluster (*Right*)*PSD(*Left*) | 0.813 | 0.393 | **2.068** |
| Condition (*Perception*)* Cluster (*Mid-right*)*PSD (*Left*) | 1.529 | 0.395 | **3.870** |
| Condition (*Perception*)* Cluster (*Centre-left*)*PSD (*Left*) | -0.969 | 0.393 | **-2.463** |
| Condition (*Perception*)* Cluster (*Left*)*PSD (*Left*) | -0.399 | 0.393 | -1.015 |
| Condition (*Perception)**Cluster(*Mid-left*)*PSD(*Left*) | -1.936 | 0.394 | **-4.918** |

#### Signed Errors Analysis

Signed errors were used to investigate if Condition, Start Position and Perspective Shift Direction (PSD) have an effect on the direction of the errors. Negative errors indicate errors to the left and positive errors are errors to the left.

Table 2 Coefficients from Signed Errors (cm) LMM analysis

|  | **Signed Errors (cm)** | | |
| --- | --- | --- | --- |
| *Predictors* | *Estimates* | *std. Error* | *t-value* |
| (Intercept) | 3.355 | 1.480 | **2.267** |
| Condition (*Memory-Perception*) | 0.815 | 1.181 | 0.690 |
| Cluster (*Right*) | -15.594 | 2.081 | **-7.493** |
| Cluster (*Mid-right*) | -4.406 | 2.082 | **-2.116** |
| Cluster(*Centre-left*) | 0.111 | 2.082 | 0.053 |
| Cluster(*Left*) | 13.861 | 2.082 | **6.658** |
| Cluster(*Mid-left*) | 3.140 | 2.082 | 1.508 |
| PSD (*Right*-*Left*) | -10.944 | 1.796 | **-6.093** |
| Condition (*Memory*-*Perception*)* Cluster (*Right*) | 0.748 | 0.595 | 1.257 |
| Condition (*Memory-* *Perception*)* Cluster (*Mid-right*) | -3.206 | 0.598 | **-5.360** |
| Condition (*Memory-* *Perception*)* Cluster(*Centre-left*) | -0.269 | 0.596 | -0.452 |
| Condition (*Memory-Perception*)* Cluster(*Left*) | -0.091 | 0.595 | -0.153 |
| Condition (*Memory-Perception*)* Cluster(*Mid-left*) | 4.614 | 0.596 | **7.741** |
| Condition (*Memory-Perception*)* PSD (*Right*-*Left*) | 0.569 | 1.559 | 0.365 |
| Cluster (*Right*)*PSD (*Right*-*Left*) | 3.942 | 2.081 | 1.894 |
| Cluster (*Mid-right*)*PSD (*Right*-*Left*) | -1.412 | 2.082 | -0.678 |
| Cluster(*Centre-left*)*PSD (*Right*-*Left*) | -2.342 | 2.082 | -1.125 |
| Cluster(*Left*)*PSD (*Right*-*Left*) | -5.245 | 2.082 | **-2.519** |
| Cluster(*Mid-left*)*PSD (*Right*-*Left*) | -1.875 | 2.082 | -0.901 |
| Condition (*Memory-Perception*)* Cluster (*Right*)*PSD (*Right-Left*) | -0.897 | 0.595 | -1.508 |
| Condition (*Memory-Perception*)* Cluster (*Mid-right*)*PSD (*Right-Left*) | 1.434 | 0.598 | **2.397** |
| Condition (*Memory-Perception*)* Cluster(*Centre-left*)*PSD (*Right-Left*) | -0.629 | 0.596 | -1.056 |
| Condition (*Memory-Perception*)* Cluster(*Left*)*PSD (*Right-Left*) | -0.354 | 0.595 | -0.594 |
| Condition (*Memory-Perception)**Cluster(*Mid-left*)* PSD (*Right-Left*) | -1.436 | 0.596 | **-2.409** |
